# Reduced frontal white matter microstructure in healthy older adults with low tactile recognition performance

**DOI:** 10.1101/775502

**Authors:** Focko L. Higgen, Hanna Braaß, Robert Schulz, Gui Xue, Christian Gerloff

**Author notes:** Corresponding author: Focko L. Higgen, University Medical Center Hamburg-Eppendorf, Martinistraße 52, 20246 Hamburg, Germany, Facsimile. Both authors contributed equally (Focko L. Higgen, Hanna Braaß).

## Abstract

Aging leads to a reduction of connectivity in large-scale structural brain networks. Sensory processing and other cognitive processes rely on information flow between distant brain areas. However, data linking age-related structural brain alterations to cognitive functioning, especially sensory processing, is sparse.

Aiming to determine group differences in sensory processing between older and younger participants, we implemented a complex tactile recognition task and investigated to what extent changes in microstructural white matter integrity of large-scale brain networks might reflect success in task performance. Structural brain integrity was accessed by means of diffusion-weighted imaging and fractional anisotrophy.

The data revealed that poor performance in complex tactile recognition in older, neurologically healthy individuals is related to decreased structural integrity pronounced in the anterior corpus callosum. This region was strongly connected to the prefrontal cortex. Our data suggests decreased fractional anisotrophy in the anterior corpus callosum as a surrogate marker for progressed brain aging, leading to disturbances in networks relevant for higher-order cognitive processing. Complex tactile recognition might be a sensitive marker for identifying these starting cognitive impairments in older adults.

## 1 Introduction

Older adults face the challenges of aging-related cognitive impairments. As some of the key features, these comprise processing of sensory stimuli, decision making and subsequent actions (Anguera and Gazzaley, 2012; Gazzaley et al., 2005; Guerreiro et al., 2014; Zheng et al., 2018). Age-related sensory impairments affect all senses, e.g. visual acuity and auditory and tactile thresholds, and have great impact on the independency and the activities of daily living (Freiherr et al., 2013). Therefore, the investigation of sensory processing might give valuable insight into mechanisms of aging.

The underlying reasons for age-related deficits are manifold. Going clearly beyond the importance of decreases of function of peripheral sensory organs, the brain undergoes continuous modifications throughout the life span (Gazzaley et al., 2005; Heise et al., 2014, 2013). These age-related alterations in central processing can be analyzed on different spatial scales. On a micro-scale, there are changes of cellular properties, morphology, transmitter levels and neural plasticity (Hong and Rebec, 2012; Kumar and Foster, 2007). On a meso-scale, these changes lead to alterations of functional neuronal activations (Heise et al., 2014, 2013; Quandt et al., 2016; Sailer et al., 2000) and consequently to large-scale alterations of brain structure and function on the network-level (Babaeeghazvini et al., 2018; Heuninckx et al., 2008; Michely et al., 2018; Raz and Rodrigue, 2006; Schulz et al., 2014).

Sensory processing relies on local brain activation and interregional information flow between primary and higher order sensory areas. Especially processing of complex stimuli needs an interplay between different brain regions and therefore requires proper structural integrity of large-scale networks (Göschl et al., 2015; Hipp et al., 2011). For tactile recognition, there is an interaction of bottom-up sensory flow with top-down control (Adhikari et al., 2014; Sathian, 2016; Stilla et al., 2007). Bottom-up tactile inputs are primarily processed in the primary somatosensory cortex (S1) and then segregated into different pathways for different object properties. Relevant cortical areas of this distributed network include the parietal operculum (SII), the posterior parietal cortices, the intraparietal sulcus, the temporo-parietal junction and the limbic areas (Mauguière et al., 1997; Sathian, 2016; Van Boven et al., 2005). Concurrently, tactile processing is mediated by higher cognitive functioning such as top-down attentional control and visuo-spatial working memory. These functions are executed via inputs from the prefrontal cortices (PFC), comprising for example the dorsolateral prefrontal cortex (DFPLC) and the ventrolateral prefrontal cortex (VLPFC) (Adhikari et al., 2014; Deibert et al., 1999; Reed et al., 2004; Sathian, 2016). Taken together, alterations in both, networks of bottom-up sensory flow and networks of top-down modulation, could lead to disturbances in sensory processing.

There have been multiple neuroimaging studies accessing large-scale brain network integrity in relation to aging processes. A measurement widely accepted to describe brain network integrity is fractional anisotrophy (FA), a parameter derived from diffusion-weighted brain imaging. Within the limitations of fiber-tracking, FA is commonly referred to as a neuroimaging index of micro-structural white matter integrity (Hugenschmidt et al., 2008; Kochunov et al., 2012; Schulz et al., 2015). In the following we will use the term accordingly.

A common finding is a wide-spread reduction of FA with increasing age (Abe et al., 2002; Carmichael and Lockhart, 2012; Kochunov et al., 2007; Madden et al., 2012, 2009; Malloy et al., 2007; Minati et al., 2007; Moseley, 2002; Salat, 2011; Sullivan and Pfefferbaum, 2007, 2006; Wozniak and Lim, 2006). This so called ‘cortical disconnection’ is thought to contribute to age-related cognitive decline (Bennett and Madden, 2014). Domains that have been shown to be affected by the decrease of micro-structural white matter integrity mainly comprise higher cognitive functioning such as executive functioning, processing speed and memory (for review see Bennett and Madden, 2014). So far, data linking these age-related structural alterations with basic sensory processing, is sparse (Chalavi et al., 2018; Damoiseaux, 2017).

Aiming to determine group differences in sensory processing between older and younger participants, we implemented a complex tactile recognition task. All participants underwent structural brain imaging including diffusion weighted imaging to characterize global white-matter microstructure. We hypothesize that inter-subject variability in microstructure of large-scale structural brain networks is related to variable success in complex sensory processing at the behavioral level.

## 2 Material and Methods

### 2.1 Participants

37 and 22 younger volunteers were screened for the study. 6 older volunteers did not meet the inclusion criteria during initial assessment. 2 older and 2 younger participants dropped out because of personal or technical problems. During task performance, 10 older participants did not meet the predefined accuracy targets (as described below) and older participants were regrouped into O-LP (older-low-performers) and O-HP (older-high-performers). Thus, 10 O-LP (5 female, mean age 74.1, range 68-82), 19 O-HP (11 female, mean age 71.9, range 65-79) and 20 younger participants (= Young, Y; 11 female, mean age 24.1, range 20-28) entered in the final analyses. All participants were right handed according to the Edinburgh handedness inventory (Oldfield, 1971), had normal or corrected to normal vision, no history or symptoms of neuro-psychiatric disorders (MMSE ≥ 28, DemTect ≥ 13) and no history of centrally acting drug intake. All participants received monetary compensation.

### 2.2 Ethics statement

The study was conducted in accordance with the Declaration of Helsinki and was approved by the local ethics committee of the Medical Association of Hamburg (PV5085). All participants gave written informed consent.

### 2.3 Assessment

Prior to inclusion, each participant underwent an assessment procedure. Assessment consisted of a neurological examination, the Mini-Mental State (MMSE, cut-off ≥ 28; Folstein et al., 1975) and the DemTect (cut-off ≥ 13; Kalbe et al., 2004) to rule out symptoms of neuro-psychatric disorders. Furthermore, a 2-point-discrimination test (cut-off > 3mm; Crosby and Dellon, 1989; Dellon et al., 1995) and a test of the mechanical detection threshold (MDT, v. Frey Filaments, OptiHair2-Set, Marstock Nervtest, Germany, cut-off > 0.75mN; Fruhstorfer et al., 2001; Rolke et al., 2006) were conducted to ensure intact peripheral somatosensation. We also assessed subjectively experienced attention deficits with a standardized questionnaire (FEDA). The FEDA is divided into sub-sections A, B and C, where A asks for distractibility and slowing up in mental processes, B for fatigue and slowing up in practical activities and C for reduction of energy (Zimmerman and Lahav, 2012).

### 2.4 Task design

The experiment took place in a light attenuated chamber. We chose experimental procedure, stimulus configuration, and stimulation parameters based on pilot data showing accuracy of tactile pattern recognition to be very different between older and younger participants.

For tactile stimulation, the participants’ right hand was resting on a custom-made board containing a Braille stimulator (QuaeroSys Medical Devices, Schotten, Germany), with the fingertip of the right index finger placed above the stimulating unit (see suppl. figure 1). The Braille stimulator consists of eight pins arranged in a four-by-two matrix, each 1mm in diameter with a spacing of 2.5mm. Each pin is controllable independently. Pins can be elevated (maximum amplitude 1.5mm) for any period to form different patterns. At the end of each pattern presentation, all pins return to baseline. The tactile recognition task consisted of steps of increasing complexity. At the beginning of each step participants read the task instructions on a computer screen positioned in front of them. The stimuli consisted of different sets of four geometric patterns, each of them formed by four dots (figure 1a). The experiment began with a very simple set of four familiarization patterns (step 1, figure 1a, b) at maximum pin amplitude and a stimulation time of 800ms to get the participants acquainted with the tactile stimulation.

**Fig. 1:**
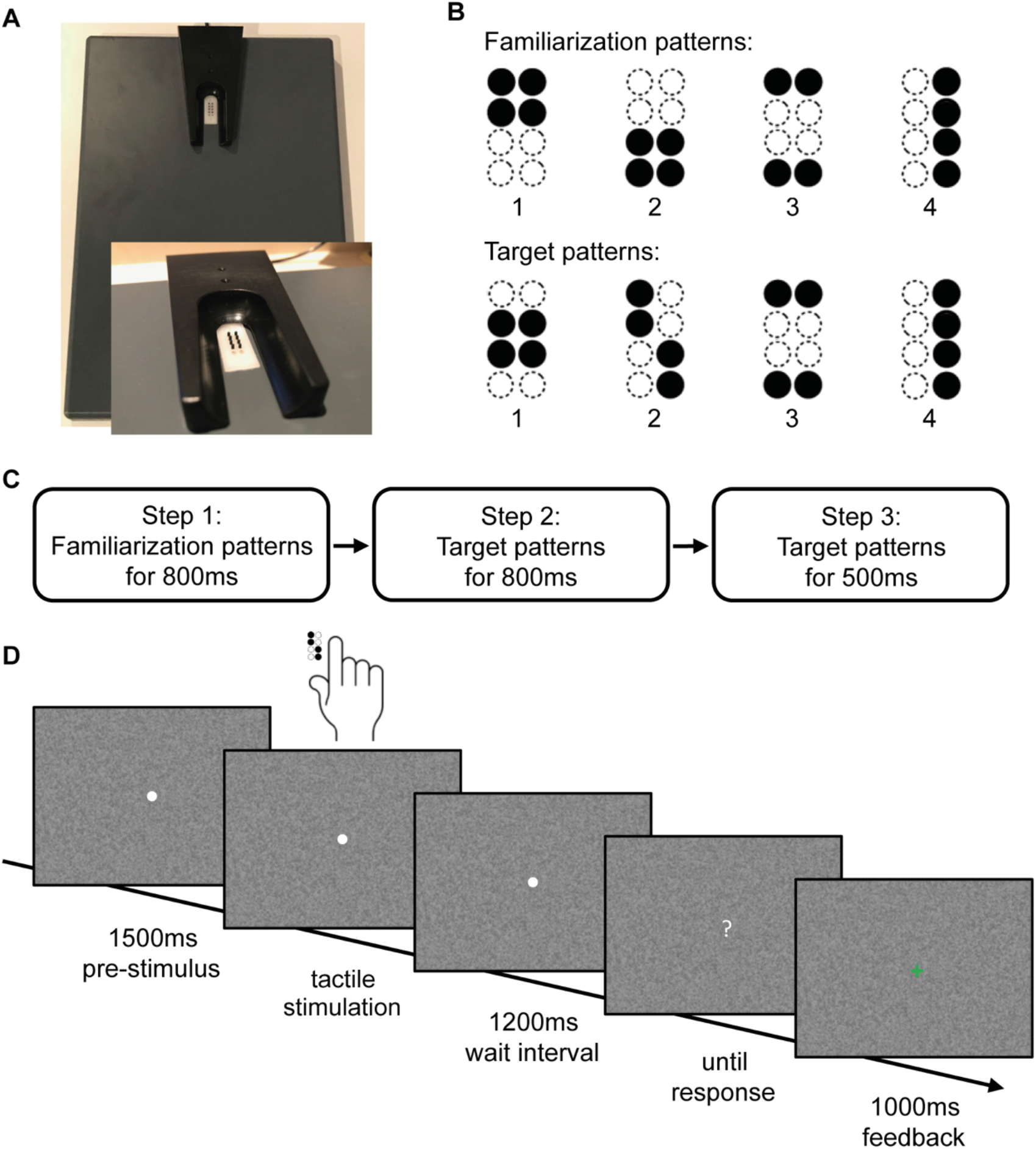
Stimulus design and experimental procedure. **A:** Braille stimulator. For tactile stimulation, the participants’ right hand was resting on a custom-made board containing a Braille stimulator (QuaeroSys Medical Devices, Schotten, Germany), with the fingertip of the right index finger placed above the stimulating unit. The Braille stimulator consists of eight pins arranged in a four-by-two matrix, each 1mm in diameter with a spacing of 2.5mm. Each pin is controllable independently. **B:** Stimuli consisted of two sets of four tactile patterns, **C:** Sequence of tasks in the experiment, **D:** The trial sequence. After a pre-stimulus interval of 1500ms, tactile patterns were presented to the right index finger with a duration depending on the current step of the experiment. After a wait interval of 1200ms, a question mark appeared on the screen and participants gave the response via button press. After response, every trial ended with a visual feedback (1000ms)

Each trial started with a central white fixation point appearing on a noisy background. This fixation point remained visible throughout each single trial. The tactile pattern presentation started 1500ms after appearance of the fixation point with a stimulus chosen pseudo-randomly from the stimulus set. After the tactile presentation, there was a waiting interval of 1200ms. Then, the central fixation point turned into a question mark and participants indicated which of the four patterns had been presented. Participants responded via button press with the fingers two to five of the left hand. After each trial participants received visual feedback (1000ms) whether response was correct (green ‘+’) or incorrect (red ‘-’) (figure 1c).

After a minimum of five familiarization blocks, each one consisting of 16 trials, and an accuracy of at least 75% in three of five consecutive blocks, participants could proceed to the next step. If participants did not reach the target accuracy within 15 blocks, they were excluded from further participation.

In the next step of the recognition task, the stimulus set consisted of the four target patterns (step 2, figure 1a, b). To train participants in the recognition of the target patterns, stimulation occurred at maximum amplitude and again with a long stimulation time of 800ms. Trial timing, blocks and accuracy targets were always as described above. If again participants were able to recognize patterns with the previously defined accuracy, in the final step of the recognition task stimulation time of the target patterns was 500ms (step 3, figure 1a, b). Participants who were able to recognize these patterns with the targeted accuracy were categorized as “high-performers”. Participants not reaching this level were labeled “low-performers”. We grouped all participants not reaching the predefined accuracy target at one of the steps of the tactile recognition task together, as they all did not show any deficits in the initial assessment, but a clear performance difference in tactile recognition compared to the younger participants and the older “high-performers”. In all these participants alterations in microstructure of large-scale structural brain networks might be the reason for poor performance, according to our initial hypotheses.

### 2.5 Brain Imaging

A 3 Tesla MRI system (Magnetom Skyra, Siemens Healthcare, Erlangen, Germany) and a 32-channel head coil acquired diffusion-weighted and high-resolution T1-weighted structural images. For the diffusion-weighted sequence, a spin-echo, echo-planar imaging (EPI) sequence was applied with the following parameters: TE = 82ms, TR = 10000ms, flip angle = 90°,matrix size = 104 × 128 matrices, FOV = 208 × 256 mm², voxel resolution = 2.0 × 2.0 x 2.0 mm³, partial Fourier factor = 0.75, 75 contiguous transversal slices, one image with b = 0 s/mm², 64 images with b = 1500s/mm² (64 non-collinear directions). For the T1-weighted sequence, a 3-dimensional magnetization-prepared rapid gradient echo (3D-MPRAGE) sequence was used with the following parameters: TR = 2500ms, TE = 2.12ms, TI = 1100ms, flip angle 9°, 256 coronal slices with a voxel size of 0.83 x 0.94 x 0.83mm³, FOV = 240 mm.

### 2.6 Image processing

The processing and analysis of MRI data were carried out using FMRIB Software Library (FSL) software 5.0.2.2 (Analysis Group, FMRIB, Oxford, UK Oxford Centre for Functional Magnetic Resonance Imaging of the Brain Software Library, https://fsl.fmrib.ox.ac.uk/fsl/fslwiki/FSL) (Smith et al., 2004). First, the eddy current distortion and simple head motion of raw diffusion data were corrected, using Eddy current correction from the FMRIB’s Diffusion Toolbox (FDT) 3.0. Then, the Brain Extraction Tool (BET) v2.1 of FSL was used for brain extraction (Smith, 2002). FA images were created by fitting a tensor model in each voxel to the raw diffusion data using FDT, additionally the eigenvalue images for L1, L2 and L3 were created in the same way.

### 2.7 Tract-Based Spatial Statistics (TBSS)

Voxelwise statistical analysis of the created FA data was carried out using TBSS (Tract-Based Spatial Statistics) (Smith et al., 2006). All subjects’ FA data were nonlinearly registered to the FMRIB58-FA standard-space template (FMRIB Centre University of Oxford, Department of Clinical Neurology, John Radcliffe Hospital Headington, Oxford, UK; https://fsl.fmrib.ox.ac.uk/fsl/fslwiki/FMRIB58_FA) and aligned to the Montreal Neurological Institute (MNI) space using the nonlinear registration tool FNIRT (Andersson et al., 2007a, 2007b) as part of TBSS, which uses a b-spline representation of the registration warp field (Rueckert et al., 1999). Next, the mean FA image was created and thinned to create a mean FA skeleton, which represents the centers of all tracts common to the group. Each subject’s aligned FA data was then projected onto this skeleton and the resulting data fed into voxelwise cross-subject statistics. The permutation-based non-parametric inferences within the framework of the general linear model were performed to investigate the differences between the groups Young versus older-high-performers (Y vs. O-HP), young versus older-low-performers (Y vs. O-LP) [1 - 1; −1 1] and older-high-performers (O-HP) versus older-low-performers (O-LP) [1 −1; −1 1] using randomise (https://fsl.fmrib.ox.ac.uk/fsl/fslwiki/Randomise). For corrected results the threshold-free cluster enhancement with the family-wise error (FWE) correction for multiple comparisons corrections (P<0.05, FWE corrected, 5000 permutations) was used.

The brain regions in which the analysis showed significantly different FA-values between the two groups were thresholded at P-value <0.05. In the next step we used these resulting brain regions to create individual masks for each subject. To create these individual masks the resulting brain regions in the anterior corpus callosum, were first binarized and in a second step back-transformed to the individual diffusion space. The individual binary masks were then multiplied with the individual FA-maps. We calculated the mean FA in the resulting image for each participant and used the mean FA-values for further statistical calculations. Mean FA values measured in a second region in the splenium corpus callosum in all older participants served as a control. The region in the splenium was manually defined on the MNI152_T1_1mm image in the Splenium corpus callosum, symmetrical to the midline of the brain (volume: 229 voxel). Additionally, axial (AD) and radial diffusivities (RD) in the anterior corpus callosum were computed in the same way.

### 2.8 Probabilistic Tractography

After preprocessing the DWI-data with eddycorrect and BET as described above FSL’s bedpostx was used to estimate the distribution of diffusion parameters in each voxel, modelling crossing fibers using Markov Chain Monte Carlo sampling (Behrens et al., 2007). Probabilistic tractography was used to reconstruct the tracts with the region defined in the anterior corpus callosum (as mentioned above) as seed mask (5000 streamlines sent from each voxel in the individual seed masks, curvature threshold 0.2, steplength 0.5mm). In each participant, tracts starting from the defined seed mask in the anterior corpus callosum were reconstructed. Group- and tract-specific connectivity distributions were finally analyzed applying different thresholds, 0.01%, 0.5%, 1.0% and 2%, of the overall successful streamlines as described elsewhere (Schulz et al., 2015).

### 2.9 Further Statistical Analyses

Further statistical analyzes were performed using Matlab version 9.1 (R2016b, MathWorks, Natick, MA) and R statistical package Version 3.5.4 (http://www.r-project.org/).

To test for assessment related group-differences a linear model was defined by means of R’s *lm* command to investigate the relationship between the assessment variables Age, MDT, 2-point-discrimination, MMSE, DemTect, FEDA-A, FEDA-B, FEDA-C as dependent variables and GROUP (Young, O-HP, O-LP) as independent variable. Age was included into the model to test for age differences in the groups O-HP and O-LP. The comparison was performed using *lsmeans* (R-package: lsmeans) and pairwise comparison between the resulting contrasts. Benjamini-Yekutieli adaptive FDR-correction (BY) was used to adjust for multiple comparisons (Benjamini and Yekutieli, 2001). For post-hoc testing a MANOVA was used with GROUP as independent variable and BY correction to adjust for multiple comparisons. Task performance of groups Young and O-HP was compared at each step of the recognition task with a two-sided t-test and BY correction for multiple comparison.

Mean FA-values in anterior and posterior corpus callosum were compared between O-HP and O-LP using a two-sided unpaired t-test. For the comparison of AD and RD in the anterior and posterior corpus callosum between the two groups O-HP and O-LP a two-sided unpaired t-test was used likewise.

Furthermore, linear regression models were fitted to test for relationships between diffusion parameters (FA, AD and RD) and assessment parameters. Group differences were calculated by means of (*diffusion parameter*)*GROUP. FDR-correction was performed to correct for multiple comparisons.

## 3 Results

### 3.1 Behavior

At all steps of the tactile recognition task, younger participants (= Young, Y) performed better than older participants (p < 0.001 at all steps). Besides this expected performance differences between younger and older participants, there were also differences in the accuracy of tactile pattern recognition within the older group. In the older group, 19 of 29 participants were able to reach the predefined accuracy level at all steps. On each step of the tactile recognition task, some older participants failed to reach the predefined target. 5 older participants were not able to detect the familiarization patterns with the targeted accuracy. 3 older participants failed to detect the target patterns at a stimulation time of 800ms. 2 more older participants failed to detect the target patterns at a stimulation time of 500ms. These participants were then excluded from the further steps. Older participants were regrouped according to their performance in O-HP (older-high-performers) and O-LP (older-low-performers). Taking only the O-HP, the younger participants still performed significantly better at each step (see table 1, p < 0.001 at all steps).

**Tab.1:** Performance of the different groups.

| Task | Young (n=20) | O-HP (n=19) | O-LP (n=10) |
| --- | --- | --- | --- |
| Familiarization<br>patterns for 800ms | 96.9 (± 3.7)* | 85.2 (± 9.7)* | 47.8 (± 8.0)<br>(n=5) |
| Target patterns<br>for 800ms | 93.1 (± 5.2)* | 76.4 (± 8.9)* | 54.4 (± 1.7)<br>(n=3) |
| Target patterns<br>for 500ms | 97.7 (± 5.1)* | 82.6 (± 10.7)* | 54.4 (± 6.8)<br>(n=2) |

Mean values over all blocks needed per participant are shown in % ± standard deviation for each step of the tactile recognition task. Group comparisons between Young and O-HP were calculated at each step of the recognition task with a two-sided t-test and BY correction for multiple comparison. \* indicate significant differences between Young and O-HP, all p-values ≤ 0.001.

### 3.2 Assessment

As the factor age was included into the model, group comparison of baseline data obtained in the assessment prior to inclusion (see table 2) naturally showed significant differences between Young and O-HP (t(46) = 37.8, p < 0.001) and Young and O-LP (t(46) = 32.2, p < 0.001). Importantly, there was no difference between O-HP and O-LP (t(46) = −0.93, p = 0.6550). Despite their differences in performance in the tactile recognition task, there were no significant differences between O-HP and O-LP in the baseline data.

**Tab. 2:** Assessment data of the groups.

| Metrics | Young (n=20) | O-HP (n=19) | O-LP (n=10) |
| --- | --- | --- | --- |
| Age | 24.1 (± 2.6)*<sup>+</sup> | 71.9 (± 4.4)* | 74.1 (± 3.9)<sup>+</sup> |
| Height (m) | 1.76(± 0.09) | 1.69 (± 0.09) | 1.71 (± 0.12) |
| Weight (kg) | 71.3 (± 12.6) | 68.2 (± 11.3) | 80.8 (± 23.2) |
| BMI (kg/m<sup>2</sup>) | 22.9 (± 2.4) | 23.8 (± 2.9) | 27.1 (± 5.7) |
| Education (years) | 12.5 (± 0.6) | 10.7 (± 1.6) | 11.7 (± 2.0) |
| DemTect | 17.8 (± 0.6)* | 16.0 (± 1.6)* | 16.9 (± 1.6) |
| MMSE | 29.7 (± 0.6) | 29.5 (± 0.6) | 29.2 (± 0.8) |
| 2-Point (mm) | 2.1 (± 0.2) | 2.2 (± 0.4) | 2.4 (± 0.5) |
| MDT (mN) | 0.28 (± 0.1)<sup>+</sup> | 0.56 (± 0.5) | 0.65 (± 0.4)<sup>+</sup> |
| FEDA A | 4.28 (± 0.4) | 4.35 (± 0.4) | 4 (± 0.5) |
| FEDA B | 4.55 (± 0.4) | 4.65 (± 0.4) | 3.97 (± 0.8) |
| FEDA C | 4.35 (± 0.6) | 4.38 (± 0.5) | 3.75 (± 0.9) |

Mean values are shown ± standard deviation. Based on significant main effects, post-hoc tests were conducted. \* indicate significant differences between Young and O-HP, <sup>+</sup> indicate significant differences between Young and O-LP, all p-values ≤ 0.01

Post-hoc comparison of baseline data of Young and O-HP, showed that besides age (F(1, 37) = 1768.1, p < 0.001) DemTect (F(1, 37) = 22.2, p < 0.001) differed significantly between groups. Young and O-LP differed, besides age (F(1, 28) = 1827.6, p < 0.001), in MDT (F(1, 28) = 18.7, p = 0.0019), but not in DemTect (F(1, 28) = 4.7, p = 0.1401). Importantly, neither of the measurements revealed pathological results in the older participants. All comparisons were corrected for multiple comparison.

### 3.3 TBSS

#### 3.3.1 Young vs O-HP and Young vs O-LP

Whole brain TBSS-analysis showed significantly higher FA-values for almost every region within the FA-skeleton for the younger participants compared with O-HP and O-LP (see figure 2). No region showed higher FA-values for O-HP or O-LP compared with Young.

**Fig. 2:**
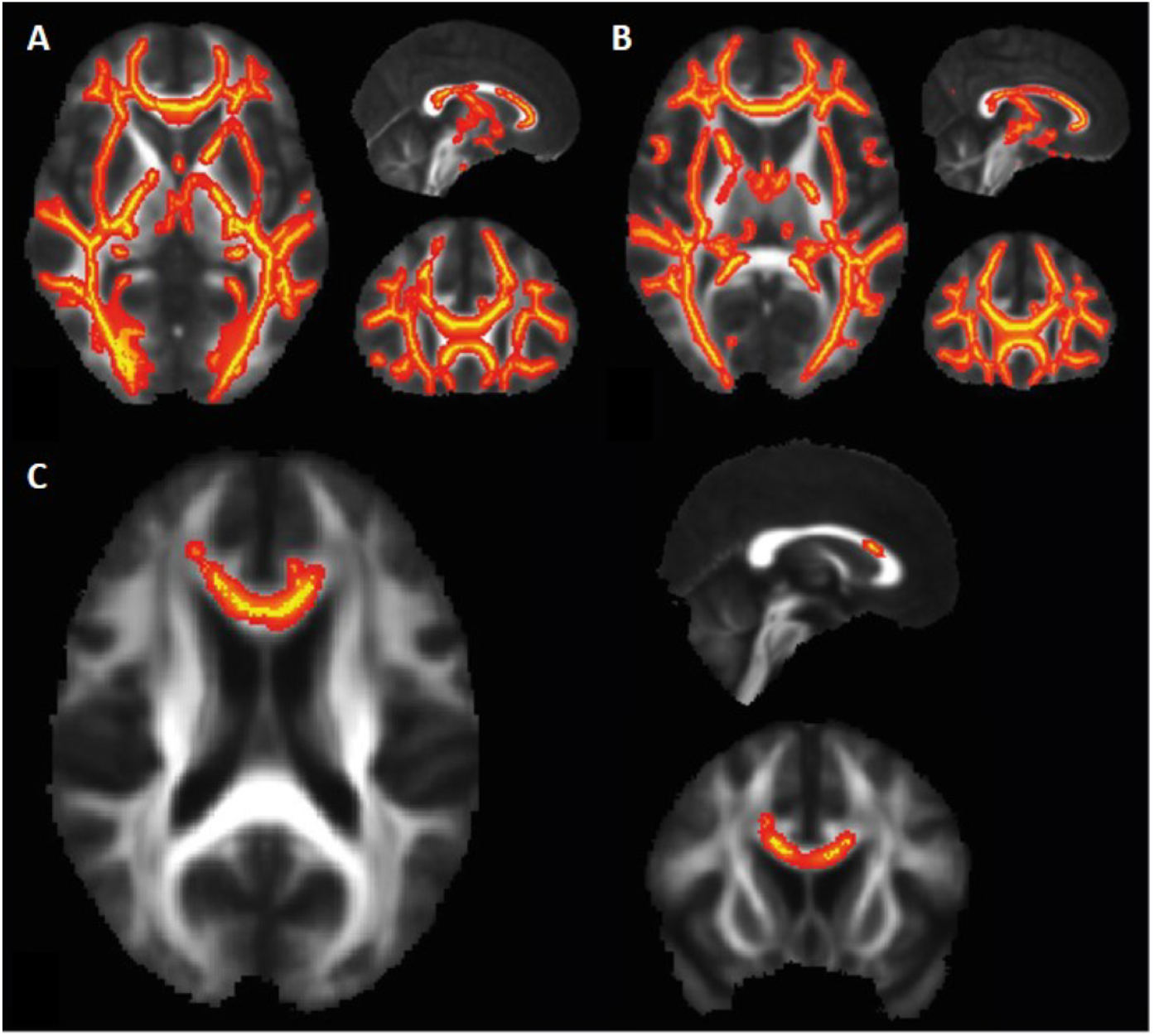
TBSS-Results, FA. **A: Y** > **O-HP**, p<0.05 (red), FWE-corrected, projected on the mean-FA, MNI-coordinates (x,y,z)=(90,153,92), **B: Y** > **O-LP**, p<0.05 (red), FWE-corrected, projected on the mean-FA, MNI-coordinates (x,y,z)=(90,155,92), **C: O-HP** > **O-LP**, p<0.05, FWE-corrected, projected on the mean-FA, MNI-coordinates (x,y,z)=(90,141,90)

#### 3.3.2 O-HP vs. O-LP

Whole brain TBSS-analysis with testing for differences between the O-HP and O-LP showed significantly higher FA-values for O-HP mainly in a region in the anterior part of the corpus callosum and a small region in the right anterior white matter connected to the corpus callosum. This part is equivalent to the overlap between genu and body of the corpus callosum (see figure 2C). The mean FA-value of this anterior region showed a significant difference between O-HP and O-LP (p < 0.001). Mean FA-values for O-HP (0.65 ± 0.02) and O-LP (0.58 ± 0.05) are additionally plotted in figure 3A. In comparison, an equivalent region in the splenium corpus callosum showed no significant difference between both groups (see suppl. figure 2).

**Fig. 3:**
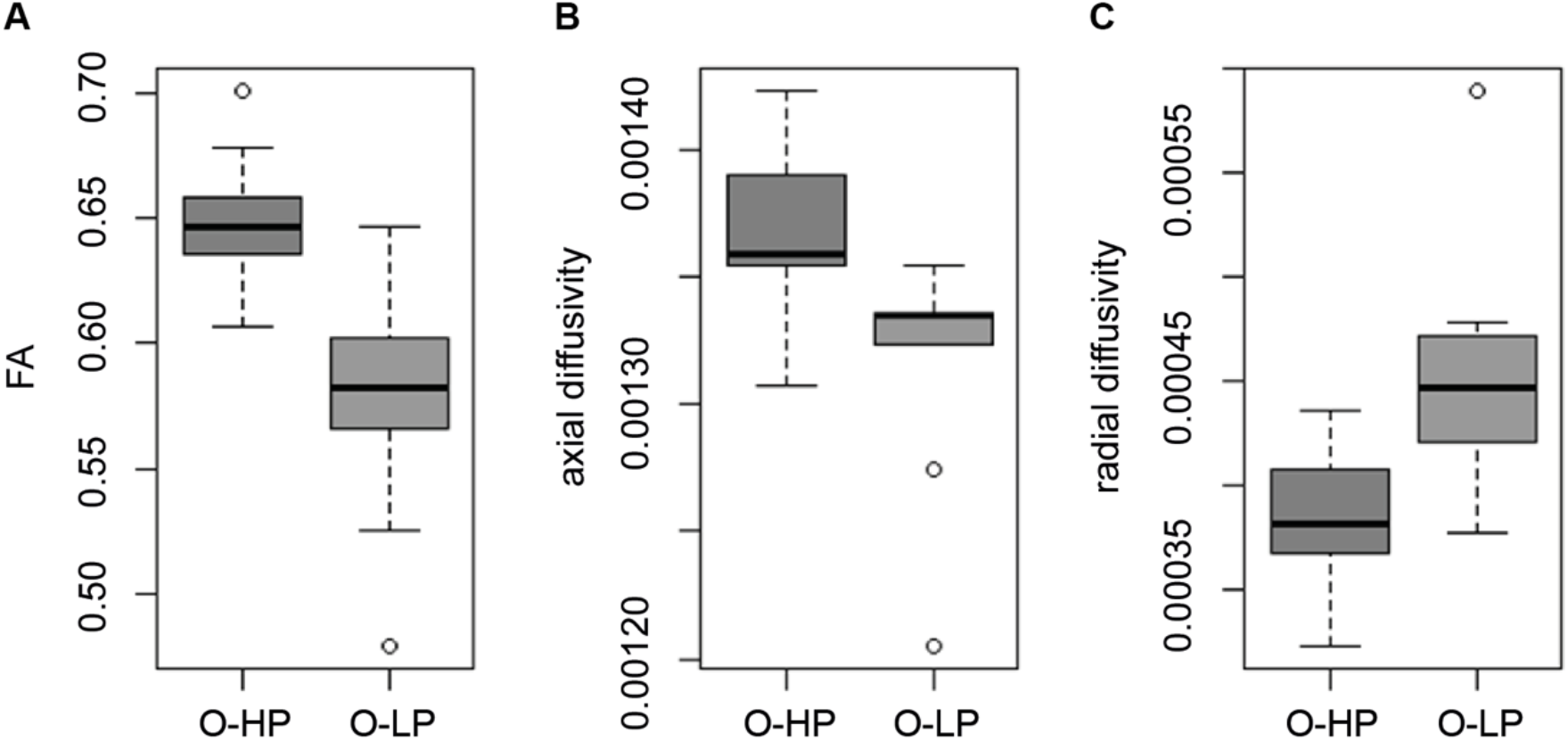
Mean FA, AD and RD for O-HP and O-LP in the anterior corpus callosum. **A:** mean FA, two-sided t-test, p<0.001 (O-HP>O-LP), **B:** mean AD, two-sided t-test, p=0.001 (O-HP > O-LP), **C:** mean RD, two-sided t-test, p<0.001 (O-LP>O-HP)

Whole brain TBSS-analysis for AD and RD showed no significant difference between both older groups within the ROI in the anterior corpus callosum. To further explore the underlying microstructural alterations in the anterior corpus callosum as defined by the TBSS-FA, we opted to investigate this region in detail. As illustrated by figure 3B and C analysis of AD showed significantly lower values for O-LP (0.00132 ± 0.00004) compared to O-HP (0.00137 ± 0.00003, p = 0.001), whereas RD was significantly higher in the group of O-LP (0.00045 ± 0.00006) compared to O-HP (0.00038 ± 0.00003, p < 0.001)

### 3.4 Tractography

To further explore the relevance of the TBSS results, we used probabilistic tractography to reconstruct the tracts originating from the region in the anterior corpus callosum. In each participant, tracts starting from the defined seed mask in the anterior corpus callosum were reconstructed.

As indicated by figure 4, there was a substantial spatial overlay of the trajectory maps for the resulting tracts for O-HP and O-LP. For each threshold, the merged tracts of all participants, thresholded by 50% of all participants of each group (O-HP and O-LP), were plotted. The reconstructed tracts showed connections between the seed-roi and the frontal lobe in both hemispheres, comprising the bilateral frontal pole, the superior frontal, inferior frontal and middle frontal gyrus, areas which are part of the prefrontal cortex. Furthermore, there were relevant connections to subcortical structures such as the bilateral thalamus and the basal ganglia. Visual inspection did not show group differences in the connected regions.

**Fig. 4:**
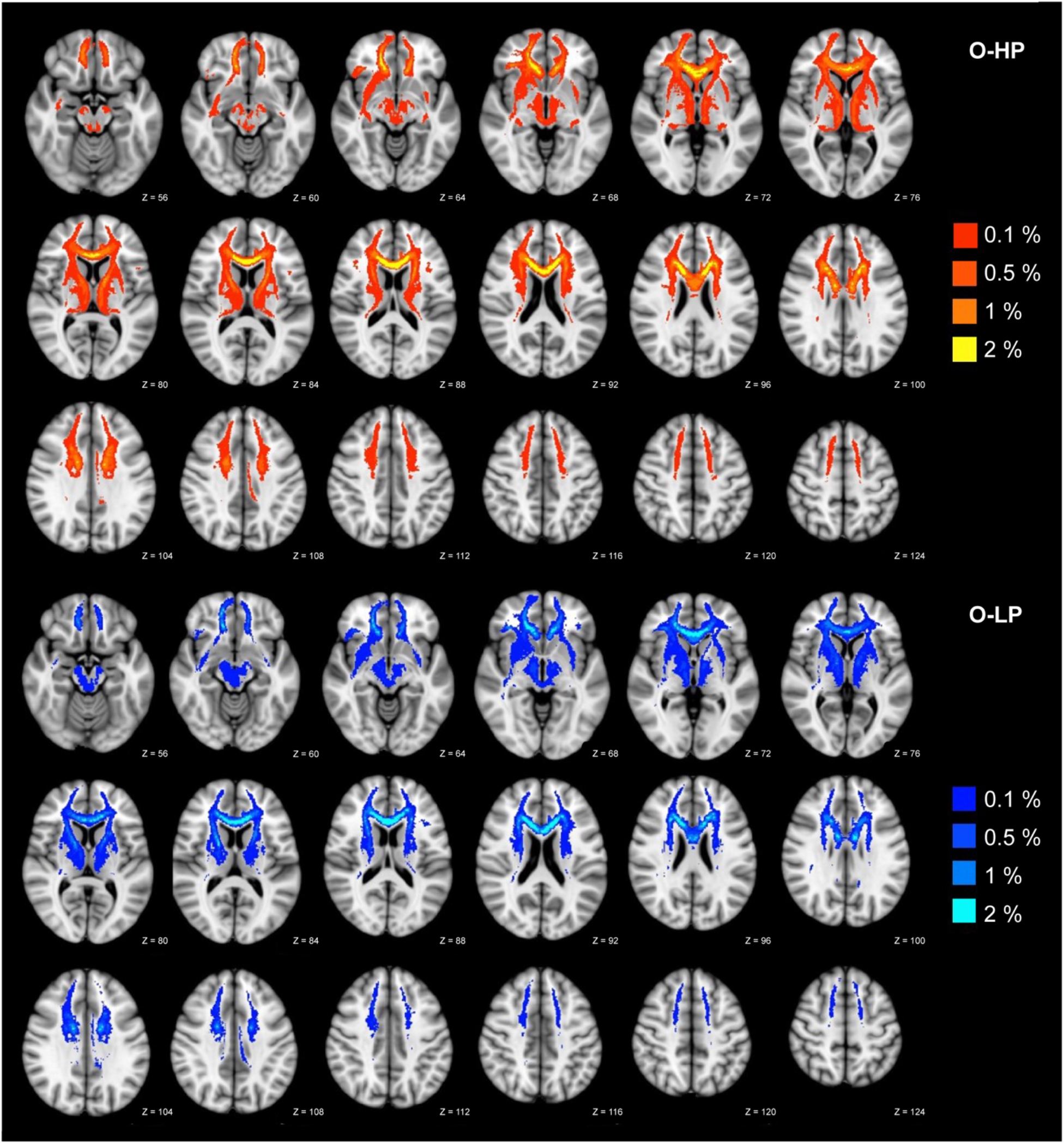
Trajectory map for the resulting tracts for O-HP and O-LP. The resulting connections are superimposed on a MNI T1 template for both groups with given z-values. Overlay of binarized group average tracts, thresholded by 50% for both groups. Individual tractography was conducted applying 5000 streamlines per voxel and thresholded by 0.1–2.0% of successful streamlines.

## 4 Discussion

This study aimed to explore complex sensory processing in younger and healthy older participants and to test the hypothesis that white matter structure has an impact on tactile behavioral performance. The data showed that, over all, older participants performed worse in a complex tactile recognition task. Intriguingly, a subgroup of the older participants showed particular low performance (O-LP), in contrast to another better performing older subgroup (O-HP). Diffusion-weighted imaging helped to better understand this bimodal distribution. The main finding was a significantly reduced microstructural integrity of transcallosal fibers, particularly in the anterior corpus callosum, in O-LP compared to O-HP. This performance-related alteration of brain structure might serve as a surrogate marker for early structural network alteration leading to differences in central processing and ultimately performance.

A decrease of FA with increasing age has been reported in multiple previous studies (Abe et al., 2002; Kochunov et al., 2007; Moseley, 2002). During aging, FA in genu und body of the corpus callosum has been shown to decrease earlier than in other regions (e.g. splenium corpus callosi) (Bennett et al., 2010; Burzynska et al., 2010; Kochunov et al., 2012; Madden et al., 2007, 2004; Michielse et al., 2010; Pfefferbaum et al., 2000; Sullivan et al., 2001). In the present data, O-HP and O-LP only differed in the FA in this area. In the context of alternative diffusion metrics, that were reduced AD and increased RD, the reduction of white-matter integrity of transcallosal fibers could be a result of age-dependent alterations both in myelinisation and axonal integrity (Alexander et al., 2011; Feldman et al., 2010).

The potential meaning of this reduced microstructural integrity in the anterior corpus callosum as a marker for early structural network alterations might be supported by its neuroanatomical properties. It has been shown that the anterior part of the corpus callosum mainly contains thinly myelinated, densely packed fibers that connect pre-frontal brain areas (Kochunov et al., 2007). These fibers maturate later and exhibit earlier deterioration during aging than the more thickly myelinated fibers in the body and splenium of the corpus callosum connecting motor or sensory areas (Bartzokis, 2004; Brickman et al., 2012; Kochunov et al., 2007). It has been hypothesized that the oligodendrocytes that myelinate the tracts passing the anterior corpus callosum are among the most metabolically active cells in the adult nervous system. This would make these cells susceptible to accumulation of metabolic damage and proposes a potential hypothesis on why the anterior corpus callosum might be more vulnerable to aging processes than other brain regions (Bartzokis, 2004; Kochunov et al., 2007). This hypothesis might provide a pathophysiological basis for our results and might indicate that reduced microstructural integrity could be caused by specific properties of the fiber system passing the anterior corpus callosum (Bennett and Madden, 2014; Salat, 2011; Salat et al., 2005).

The anterior corpus callosum has been shown to mainly connect pre-frontal brain areas (Kochunov et al., 2007) We used probabilistic tractography to further relate the local FA reduction in the anterior corpus callosum in O-LP to the underlying structural networks.

Information flow between distant brain regions has been shown to be of critical importance for processing of sensory information (Ni and Chen, 2017). As discussed in the introduction, sensory processing relies on distributed networks relevant for bottom-up sensory flow and top-down control. Alterations in both might lead to disturbances in tactile recognition. Hence, the identification of specific neuronal networks affected by the decrease in microstructural white matter integrity might give insights into the reasons for poor task performance in O-LP (Coxon et al., 2012).

Probabilistic tractography showed strong connections from the anterior corpus callosum to the frontal pole, the inferior, middle and superior frontal gyrus, heterogeneous brain regions contributing to prefrontal cortices (PFC), such as the dorsolateral (DFPLC) and ventrolateral (VLPFC) prefrontal cortex and also the orbitofrontal cortex (OFC). The PFC has been shown to be relevant for various higher-order cognitive processes, e.g. working memory (Funahashi, 2017; Miller et al., 2002; Miller and Cohen, 2001). More specifically, the DFPLC is known for its role in the executive functions, such as selective attention and cognitive flexibility (Curtis and D’Esposito, 2003; Gläscher et al., 2012; Kim et al., 2011). The VLPFC has been reported to be an important node in elaborate attentional processes and top-down processing of sensory information (Tops and Boksem, 2011; Uno et al., 2015). The OFC is involved in decision making (Fellows, 2007; Wallis, 2007). In addition to possible connections between these cortical brain regions, higher cognitive processes are also reliant on cortico-subcortical circuits, connecting cortical brain areas with the thalamus (Behrens et al., 2003; Ferguson and Gao, 2015) and the basal ganglia (McNab and Klingberg, 2008; Voytek and Knight, 2010). Well in line with this, the identified region in the anterior corpus callosum was also found to be connected bilaterally to the thalamus and the basal ganglia.

Taken together, probabilistic tractography confirmed that the identified region in the anterior corpus callosum mainly connects pre-fontal cortices. The reduction of microstructural integrity in the anterior corpus callosum might lead to functional disturbances in these frontal cortico-cortical and cortico-subcortical networks and impede cognitive processes related to attention and working-memory which are important for complex sensory information. Linking the structural finding to the behavioral results, one could argue that the disturbance of the identified networks might be a possible reason for poor task performance in O-LP. Interestingly, there were no relevant structural connectivity between the seed region and parietal brain areas, suggesting that poor performance of O-LP seems not to be primarily related to networks relevant for primary central stimulus processing.

In order to exclude potential confounding factors, we also investigated alternative reasons for the group difference between O-HP and O-LP. First, there was no age difference between O-HP and O-LP. Second, there was no difference in peripheral somatosensation between the two groups. In contrast, behavioral tests comparing younger and older individuals pointed to a difference in the MDT which tended to be worse in the older participants. Of note, this difference became only significant when comparing Young selectively to O-LP. Third, the was no difference in the neurophysiological assessment between O-HP and O-LP. O-LP even had higher scores in DemTect and did not differ significantly from Young. While all participants with pathological test results were excluded, there still was a significant difference between Young and O-HP in DemTect.

Taken together, possible reasons of performance differences between younger and older participants are manifold, comprising differences in peripheral stimuli processing, cognitive decline, but also structural changes of the brain. Comparing O-HP and O-LP, the only difference was found in brain imaging and the regional microstructure in the anterior corpus callosum. We argue that this might be a specific finding showing a progressed aging of the brain and explaining performance differences.

On a speculative note, biological age might not and neurophysiological assessment might not yet reflect the progressed aging of the brain in O-LP. The complex tactile recognition task might be a sensitive marker for a more general early cognitive impairment, potentally more sensitive than common neurophysiological measures. As one of the most important endeavors in this field is to identify individuals suffering from age-related impairments to allow for early support and interventions, complex tactile recognition might be one asset. Still, due to the found alterations in networks relevant for higher order cognitive processes, early cognitive impairment might also be assessed by complex sensory tasks in other modalities, which rely on the same frontal networks connected by the anterior corpus callosum.

There are some limitations to the current study. Due to group allocation based on performance, sample size of O-HP and O-LP differed which might limit statistical robustness of the results. Additionally, overall sample sizes were small and this might question the generalizability of the present results. Especially for any negative results we cannot preclude that a larger sample size would detect smaller effects. This is why we did not report any small effects but restricted our analyzes and interpretation to the very large effect found in the anterior corpus callosum. Though, the current study might help to generate hypotheses for further explorations. For instance, prospective studies would help to investigate structural integrity in the anterior corpus callosum over time. Likewise, the effects of behavioral training on FA in this region could be evaluated by future longitudinal studies, which thereby might also infer causality of the present findings.

In conclusion, the only difference found between healthy older low- and high performers in a complex tactile recognition task was a decreased microstructural white-matter integrity in the anterior corpus callosum. In line with the literature, showing that due to it’s neuroanatomical properties the anterior corpus callosum is most vulnerable to aging processes, we argue that this might be a specific finding showing a progressed aging of the brain and explaining performance differences. Sensory processing relies on both, primary sensory processing and higher cognitive processes. As the vulnerable region in the anterior corpus callosum mainly connects pre-frontal cortices, our results suggest that disturbances in higher order cognitive processes are the reason for early decline with aging.

